# SART1 localizes to spindle poles forming a SART1 cap and promotes spindle pole assembly

**DOI:** 10.1101/2023.10.27.564116

**Authors:** Hideki Yokoyama, Daniel Moreno-Andrés, Kaoru Takizawa, Zhenzhen Chu, Anja Scheufen, Tsumugi Funabashi, Jian Ma, Wolfram Antonin, Oliver J. Gruss, Yoshikazu Haramoto

**Affiliations:** National Institute of Technology, Ibaraki College, 312-8508 Hitachinaka, Japan; Institute of Biochemistry and Molecular Cell Biology, Medical School, RWTH Aachen University, 52074 Aachen, Germany; ID Pharma Co. Ltd., 300-2611 Tsukuba, Japan; Institute of Genetics, University of Bonn, Karlrobert-Kreiten-Str. 13, 53115 Bonn, Germany; National Institute of Advanced Industrial Science and Technology, 305-8566 Tsukuba, Japan

**Keywords:** centrosomal protein, microtubule binding, PCM proteins, spindle pole formation, Ran

## Abstract

The nuclear protein SART1 has been associated with pre-mRNA splicing but SART1 RNAi knockdown results also in defects in mitotic progression, centrosome biogenesis and chromosome cohesion. The mitotic roles of SART1 have not been characterized in detail and it remains unclear whether SART1 functions in mitosis directly or indirectly via pre-mRNA splicing. Here, we identify SART1 as a direct, mitosis-specific microtubule-associated protein. SART1 downregulation in human cells leads to spindle assembly defects with reduced microtubule dynamics, lack of end-on attachment, and checkpoint activation, while microtubule dynamics remain unaffected in interphase. SART1 uniquely localizes to the distal surface of mitotic centrosomes along the spindle axis, forming a previously not described structure we refer to as SART1 cap. Immunoprecipitation of SART1 consistently identifies centrosomal proteins as interaction partners. Immunostaining of these shows that SART1 downregulation does not affect centriole duplication and centrosome-accumulation of γ-tubulin but reduces the accumulation of selective pericentriolar material (PCM) proteins like Ninein. Depletion of SART1 from frog egg extracts disrupts spindle pole assembly around sperm nuclei and DNA-coated beads. Spindles formed around DNA-coated beads do not contain centrosomes but still recruits PCM proteins for spindle pole assembly. We finally show that the N-terminus of SART1 is its microtubule-binding region and essential for spindle assembly. Our data unravel a unique localization of SART1 and a novel function to recruit selective PCM proteins for spindle pole assembly in centrosomal and acentrosomal spindle assembly.

## Introduction

The squamous cell carcinoma antigen recognized by T-cells 1 (SART1) gene was first identified and cloned as a carcinoma antigen recognized by cytotoxic T-cells (Shichijo et al., 1998). The SART1 protein is expressed in most proliferating cells but overexpressed in some cancer cells including epithelial cancers (Cromer et al., 2004). Therefore, SART1 has been considered as a potential diagnostic marker and therapeutic target in various cancers (Allen et al., 2012; Olson et al., 2010). However, it is still unknown whether SART1 overexpression is a cause or a consequence of malignant cell transformation and the physiological function of SART1 remains incompletely understood.

In cell free assays using HeLa cell nuclear extracts, SART1 is required for U4/U6.U5 tri-snRNP recruitment to the pre-spliceosome and thus plays a role in pre-mRNA splicing (Makarova et al., 2001). Accordingly, SART1 is also called as U4/U6.U5 tri-snRNP-associated protein 1. In addition, SART1 was identified in genome wide RNAi screens as a factor required for mitotic progression (Kittler et al., 2004; Neumann et al., 2010). Specific RNAi screens identified SART1 as a critical factor for centriole biogenesis (Balestra et al., 2013) and sister chromatid cohesion (Sundaramoorthy et al., 2014). SART1 knockout in mice is embryonically lethal (Koh et al., 2016). Taken together, this suggests that SART1 has crucial functions in mitosis and cell division with vital importance for organismic homeostasis and/or development. However, we still lack detailed knowledge about the molecular action of SART1 in mitosis.

In eukaryotic cells, proteins containing nuclear localization signal (NLS) are actively imported into the nucleus. During interphase, they are retained in the nucleus and might have critical nuclear functions, such as in DNA replication, transcription, and splicing. Upon nuclear envelope breakdown at the beginning of mitosis, however, nuclear proteins become accessible to cytoplasmic proteins, including microtubules (MTs), and some play cell cycle-specific, moonlighting roles in spindle assembly and function (Somma et al., 2020). Many nuclear proteins specifically function in the vicinity of mitotic chromatin. They are spatially activated via the GTP-bound form of the Ran GTPase (RanGTP) produced locally around chromatin (Gruss, 2018). RanGTP binds to the heterodimeric nuclear transport receptor importin α/β and dissociates NLS-containing nuclear proteins from these importins. The liberated nuclear proteins are active and induce spindle assembly around chromatin (Gruss, 2018). To identify such nuclear MT regulators, we have affinity-purified proteins that contain NLS sequences and bind to MTs (MT-associated proteins (MAPs)) (Christodoulou and Yokoyama, 2015). We obtained more than 200 proteins (Yokoyama et al., 2013). The NLS-MAPs identified this way have been proven to be an excellent resource to uncover new mitotic regulators and reveal their functions (Yokoyama et al., 2014; Yokoyama et al., 2019).

We show here that one of these factors is SART1, which was so far not recognized as MAP, uniquely localizes to the most far end of spindle poles and is required for mitotic progression and spindle assembly. Our data suggest that the primary function of SART1 at spindle poles is to recruit several canonical centrosomal proteins to these sites during mitosis.

## Results

### SART1 is a bona-fide microtubule-associated protein

We have previously identified SART1 as a potential MAP (Yokoyama et al., 2013). To corroborate this finding and test whether SART1 can interact with MTs, we added taxol-stabilized MTs to HeLa nuclear extracts and recovered MTs and their interacting proteins by centrifugation. Endogenous SART1 from HeLa nuclear extracts was efficiently co-sedimented with MTs, while GAPDH was not co-sedimented, indicating a specific interaction of SART1 with MTs (Fig 1A). Addition of recombinant importin α/β complex inhibited SART1-MT interaction (Fig 1A). Inhibition was reversed by the co-addition of RanGTP, which binds to importin β and prevents the importin α/β complex from binding to NLS-sites. As previously reported, the MT binding of the MT polymerase chTOG, the human orthologue of Xenopus XMAP215, showed no inhibition by importins (Yokoyama et al., 2019). Similarly to the situation in HeLa nuclear extracts, SART1 was co-sedimented with taxol-stabilized MTs in Xenopus egg extracts (Fig S1A).

**Figure 1.**
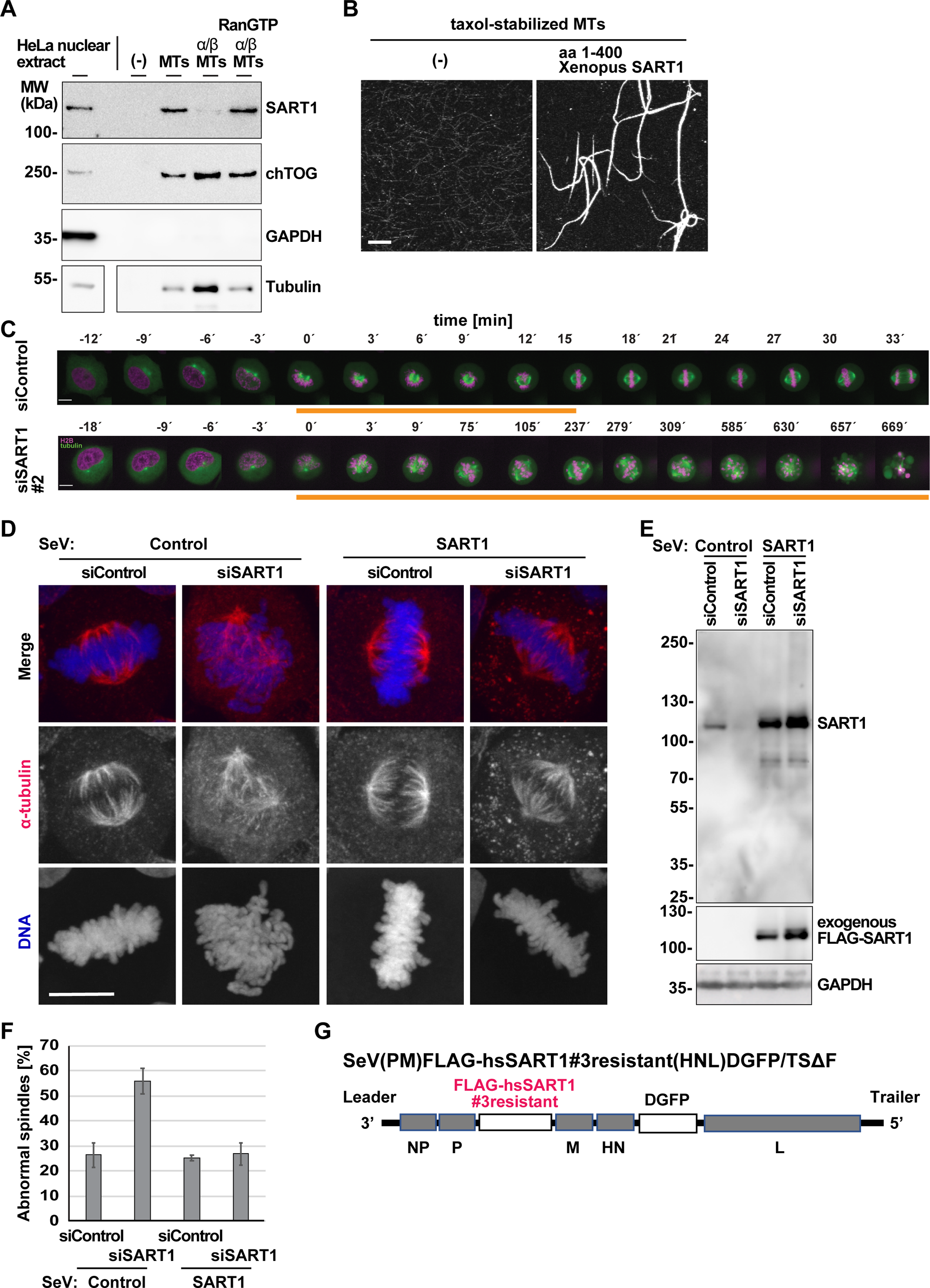
SART1 is a MAP required for early mitotic progression and spindle assembly. A. Human SART1 binds to microtubules (MTs) in a RanGTP-regulated manner. HeLa nuclear extract was incubated with taxol-stabilized MTs in the presence or absence of recombinant importin α/β complex and RanGTP and then centrifuged. Microtubule-associated proteins (MAPs) were eluted with high salt from the pellet. After centrifugation, and the supernatant (the eluate) was analyzed by western blot (WB). B. N-terminal fragment of SART1 bundles MTs in vitro. aa 1-400 Xenopus SART1 was incubated with taxol-stabilized, Cy3-labeled MTs and imaged with fluorescent microscopy. C. Confocal images HeLa cells expressing GFP-tubulin and mCherry-H2B were transfected with indicated siRNAs and imaged in intervals of three minutes. Orange lines indicate cells in prometaphase. D. SART1 is required for spindle assembly. HeLa cells were infected with control Sendai virus (SeV) vector or SeV carrying an siRNA #3-resistant version of human SART1. The cells were subsequently treated with control or SART1 siRNAs for 3 days, fixed, stained for α-tubulin and DNA, and imaged by a confocal microscope. Maximum-projected images are shown. E. Western blot of lysates from HeLa cells assayed in D. F. Quantitation of abnormal spindles assayed in D. Error bars: SD. N = 4 experiments, n > 50 prometaphase and metaphase-like cells based on chromatin shapes were counted per experiment. Note that prophase, anaphase, telophase-like cells were not considered. The abnormal spindles counted may include normal prometaphase cells going to assemble spindles later. G. Scheme of the Sendai Virus (SeV) harboring FLAG-tagged human SART1, resistant to siSART1 #3, and Dasher GFP (DGFP). Scale bars, 10 μm.

To examine whether SART1 directly interacts with MTs, we expressed recombinant Xenopus SART1 in bacteria, with an Acidic-Target Tag (Sangawa et al., 2013) for better solubility (Fig. S1B). In the MT sedimentation assay with purified components, recombinant SART1 bound to MTs (Fig. S1C), in contrast to TRIM21 which served as a negative control. A recombinant N-terminal SART1 fragment (aa 1—400) without the acidic tag bound to and bundled MTs efficiently in vitro while C-terminal fragments did not (Fig. 1B and S1D). These results indicated that SART1 is a bona-fide, previously unrecognized MAP with an N-terminal MT domain, which is regulated by importins and RanGTP.

### SART1 is required for mitotic progression and spindle assembly

To assess SART1 impact on mitosis, we knocked down SART1 in HeLa cells stably expressing tubulin-GFP and histone H2B-mCherry and performed live-cell imaging during mitosis. SART1 downregulation with three different small-interfering RNAs (siRNAs) induced persistent chromosome misalignments, spindle MT assembly defects and promethaphase-like arrest (Fig 1C and S2A). Prolonged prometaphase was confirmed by CellCognition analysis (Held et al., 2010; Yokoyama et al., 2019) (Fig S2B). Further analysis of the live cell data revealed that a considerable number of cells (15% to 45% depending the siRNA oligo) died as consequence of early mitotic defects (Fig. S3C and S3D).

Immunostaining of the fixed siSART1-treated HeLa cells showed chromosome misalignments and spindle MT defects (Fig 1D-F). MTs appeared not arranged inside the spindle, and spindle poles were not well focused. Similar spindle defects were observed with all the three siRNAs (Fig. S3A) as well as in U2OS cells (Fig. S3B). In agreement with our results from live cell imaging, we found by FACS analysis that SART1 depletion significantly induces apoptotic cell death (Figs. S3C). Among the three SART1 siRNAs, we further used siSART1#3 for detailed characterization of SART1, because it caused the lowest numbers of cell deaths.

To proof the specificity of the depletion phenotype, we performed rescue experiments. We employed a Sendai virus (SeV) vector to transiently express a SART1 version resistant to downregulation by siSART1#3 oligo (Li et al., 2000) (Fig 1G). Indeed, the exogenous SART1 was well expressed in HeLa cells and fully abolished the spindle defects caused by siSART1#3 treatment (Fig. 1 D-F).

### SART1-silencing reduces microtubule dynamics in mitosis and prevents end-on attachment of spindle microtubules to chromosomes

To understand the nature of the spindle defects, we tracked MT association and movement of EB3, a plus end MT marker, by live imaging of mitotic cells (Barenz et al., 2013). EB3 spindle/microtubule association was significantly reduced upon SART1 knockdown, suggesting a reduction of MT polymerization events (Fig. 2A and B). SART1 silencing also decreased track length and duration of MT growth while growth speed was hardly affected (Fig. 2C-E). Importantly, the reduction of track length and duration was seen for mitotic but not interphase MTs (Fig. 2C-D).

**Figure 2.**
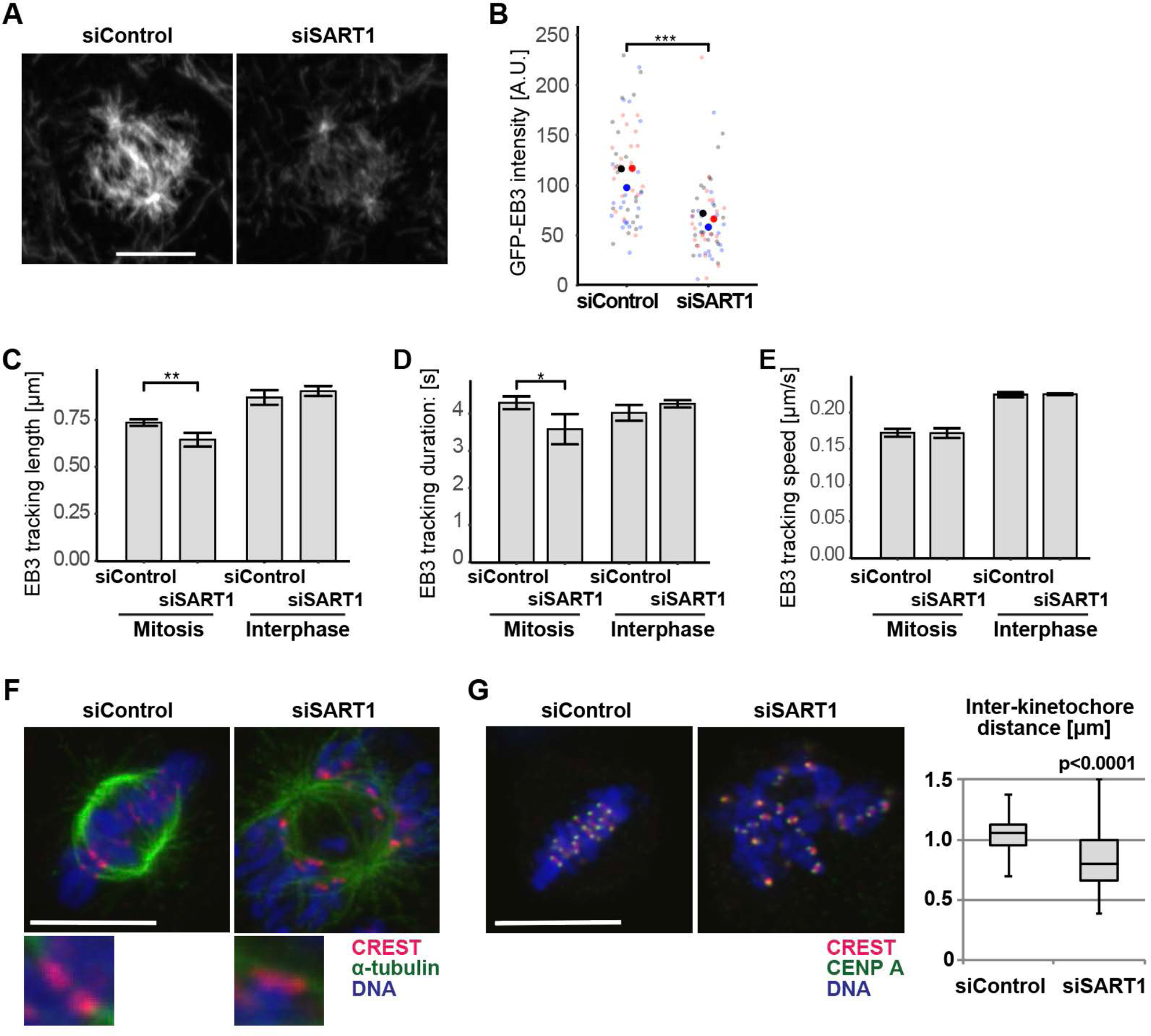
SART1-silencing reduces microtubule dynamics in mitosis and prevents end-on attachment of spindle microtubules to chromosomes. A. SART1 is required for MT plus-end binding of EB3 in mitotic MT. HeLa cells stably expressing GFP-EB3 were imaged and control cells were compared to SART3 knockdown cells. B. Reduced EB3 plus end binding in mitotic MT in the absence of SART1. EB3 intensity as the sum of all mitotic MT (see e.g. A), was quantified with ImageJ.. Overall intensity signals from 3 x 20 spindles (small dots) from 3 independent experiments (different colors, mean = large dots) are shown. The significance test (two-tailed students’ t-test) compares mean values with p≤0.001 (***). C-E. MT tracking was performed in GFP-EB3 expressing control and SART3 knockdown cells at 1 s interval for 60 s and three independent experiments with quantifications of 20 – 50 individual tracks in each of 20 spindles (i.e. >500 tracks for each condition) were performed. Mean values of these experiments are shown and compared using a two-tailed students’ t-test with p≤0.05 (*) or p≤0.01 (**). C. MT length is reduced during mitosis in the absence of SART1. D. MT growing time is reduced during mitosis in the absence of SART1. E. MT growing speed is unchanged in the absence of SART1. F. End-on MT-kinetochore interactions are impaired upon SART1 downregulation. HeLa cells were treated with the siRNAs, fixed, and stained for the kinetochore marker CREST as well as α-tubulin, and DNA. Confocal slice images are shown. Note that end-on attachment of MTs to kinetochores are seen in control cells but could not upon SART1 downregulation. 3-fold magnified images are shown below. G. Interkinetochore distance is reduced upon SART1 downregulation. The RNAi-treated cells were stained for the centromere marker CENP-A, the kinetochore marker CREST, and DNA. Confocal slices were used to find kinetochore pairs to measure interkinetochore distance. n > 94 kinetochore pairs. A two-tailed students’ t-test reveals statistical significance with p≤0.0001. Scale bars, 10 μm.

To examine the nature of chromosome misalignments, we immunostained SART1-depleted cells with human CREST autoimmune-serum, which specifically labels the kinetochore. In control cells, kinetochore pairs were connected to MTs emanating from both spindle poles suggesting end-on attachment (Fig. 2F). By contrast, in SART1 depleted cells, we could not detect such MT-kinetochore attachment (Fig. 2F). Functional end-on attachments generate tension to separate kinetochores of sister chromatids, increasing the inter-kinetochore distance (Hara and Fukagawa, 2020) and, accordingly, inter-kinetochore distances seemed to be decreased upon SART1 knock-down (Fig 2F). To confirm this, we co-stained the cells for CREST and CENP-A, an inner kinetochore marker and histone H3 variant incorporated into centromeric DNA, and measured the distance of two CENP-A dots. Whereas control cells showed an average inter-kinetochore distance of 1.1 μm, in SART1 depleted cells the distance was reduced to 0.8 μm (Fig. 2G). Staining with an outer kinetochore marker Ndc80/Hec1 also revealed a shorter interkinetochore distance upon SART1 depletion (control 1.6 μm and depletion 1.1 μm, Fig. S3D). Consistently, the mitotic checkpoint protein BubRI remained localized at the metaphase spindle (Fig. S3E), indicating the spindle assembly checkpoint (SAC) is active in the absence of SART1. Analysis of chromosome spreads showed proper sister chromatid cohesion in our depletion condition (Fig. S3F). These results indicate that SART1 is not required for sister chromatid cohesion but essential for establishing and/or maintaining MT-kinetochore attachment.

### SART1 uniquely localizes to specific surface of the centrosomes in a microtubule-dependent manner

To understand the precise subcellular localization of SART1 actions in mitosis, we examined the localization of SART1 in HeLa cells. A human SART1 antibody stained the nucleus and partially the cytoplasm in interphase cells as reported (Binder et al., 2014), but labeled the spindle poles in mitosis (Fig. 3B). The staining was lost upon SART1 downregulation and restored by expressing siRNA resistant SART1 (Fig. 3B) demonstrating its specificity. After nuclear envelope breakdown, SART1 localized around two MT asters at prophase and accumulated further around the arising spindle poles at prometaphase (Fig. S4A). The spindle pole localization was most prominent in metaphase, remained during anaphase and eventually disappeared in telophase (Fig. S4A).

**Figure 3.**
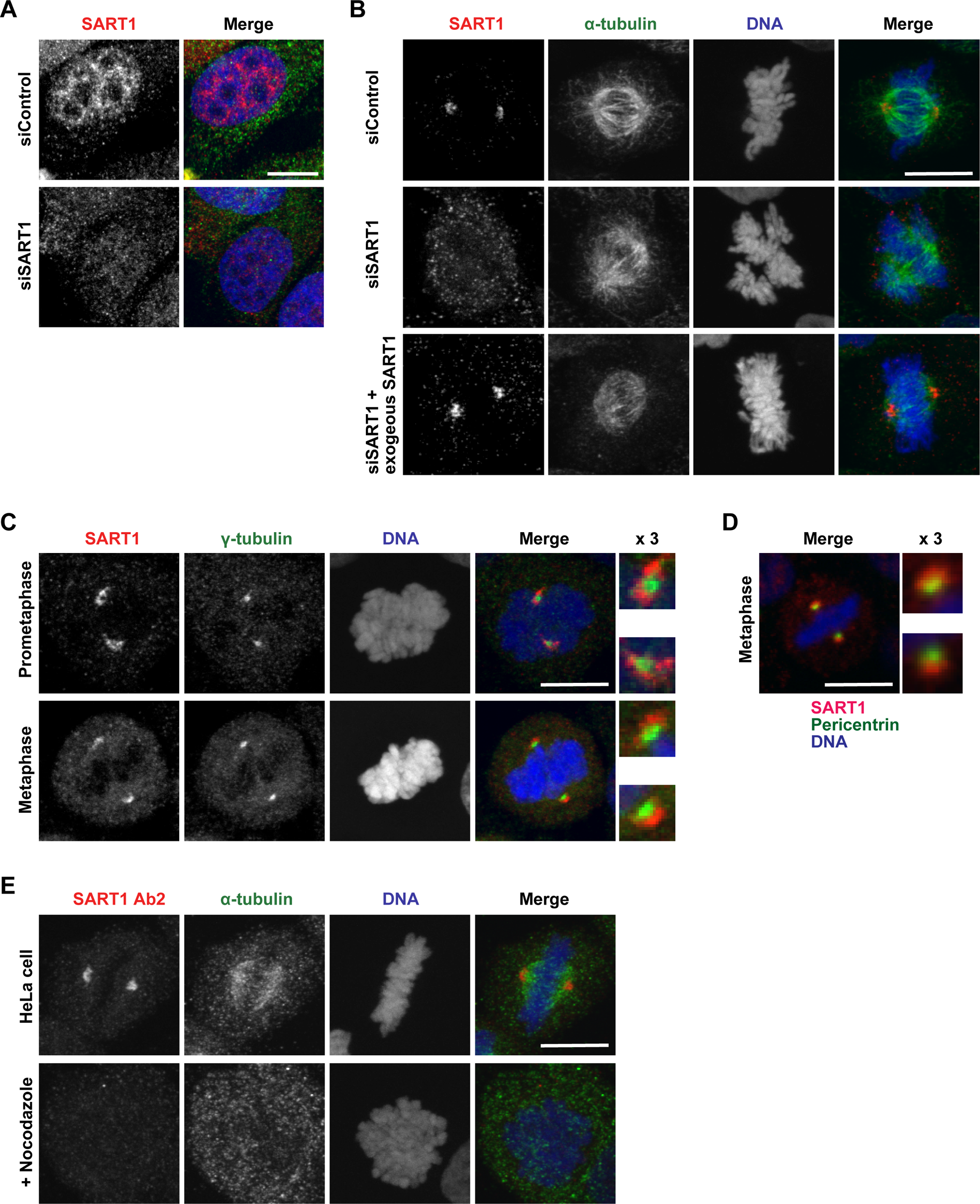
SART1 localizes to a specific subdomain of mitotic centrosomes. A and B. HeLa cells were treated with control or SART1 siRNAs for 3 days, fixed, and stained for SART1 (red), α-tubulin (green), and DNA (blue). A. In interphase, SART1 localizes within nucleus. B. SART1 localizes to spindle poles. Note that SART1 signals disappear in SART1 RNAi cells, validating the SART1 localization found in control cells. C. SART1 localizes around γ-tubulin in early mitosis, but eventually forms a half circle around the γ-tubulin distant from the spindle axis. HeLa cells were fixed, and stained for SART1, γ-tubulin, and DNA. D. SART1 localizes outside of pericentrin along the spindle axis in metaphase. HeLa cells were fixed, and stained for SART1, pericentrin, and DNA. E. The SART1 localizes at spindle poles in the presence of MTs. HeLa cells were treated with nocodazole for 10 min before fixation and stained for SART1 α-tubulin, and DNA. Scale bars, 10 μm.

The major determinant of spindle poles in somatic cells are centrosomes that function as the primary MT-organizing center. Each centrosome consists of two centrioles surrounded by the pericentriolar material (PCM) (Buhler and Stolz, 2022). Co-staining with the PCM marker γ-tubulin revealed that SART1 localized around γ-tubulin in early mitosis, but only covered the distal surface of the γ-tubulin along the spindle axis (Fig. 3C). The SART1 signal was similarly detected around pericentrin, another PCM marker (Fig. 3D). The same SART1-pericentrin configuration was also observed using a different SART1 antibody (Fig. S4B).

As we have identified SART1 as a MAP, we examined how MTs affect the spindle pole localization of SART1. Before fixation, HeLa cells were treated for 10 min with nocodazole to depolymerize MTs. In this condition, SART1 disappeared from the spindle poles (Fig. 3E), indicating a requirement of MTs for the SART1 localization. This contrasts to the pole localization of γ-tubulin which remained largely unchanged upon nocodazole treatment (Fig. S4C).

These results indicate that SART1 accumulates around centrosomes during early mitosis and, once the spindle establishes, localizes to the distal surface of centrosomes along the spindle axis. This localization is dependent on MTs.

### SART1 interacts with centrosomal proteins and recruits selective PCM proteins for spindle pole assembly

To understand the mechanism of how SART1 localizes to centrosomes and how SART1 functions there, we immunoprecipitated SART1 from Xenopus egg extracts using a specific antibody that we generated (Fig. 4A and S5A). SART1-interacting proteins were eluted from the antibody beads with a high pH buffer (Fig. S5B). These conditions revealed SART1 interaction partners while SART1 itself remained mostly on the beads (Fig. S5B) presumably due to strong antigen-antibody interaction. Analysis of the eluate by shotgun mass spectrometry identified known interacting partners of SART1 involved in mRNA splicing (SF3B1, SF3B2, IK, SF3B4, RBM25, SF3B5, PRPF40A, CHERP, DDX42, ZMAT2, SNRNP70, PRPF6, SF3A3, SNRPA, DDX46; Table S1) (Szklarczyk et al., 2021). As expected, SART1 itself was found with low mass spectrometry score. Intriguingly, mass spectrometry identified numerous centrosomal proteins (Cep192, APC, POC1, Cep44, Cep43, Cep152, Cep63, Plk1, SSX2IP, Cep97, Cep85, POC5, CDK11B, Cep70, Centrin 3; Table S1).

**Figure 4.**
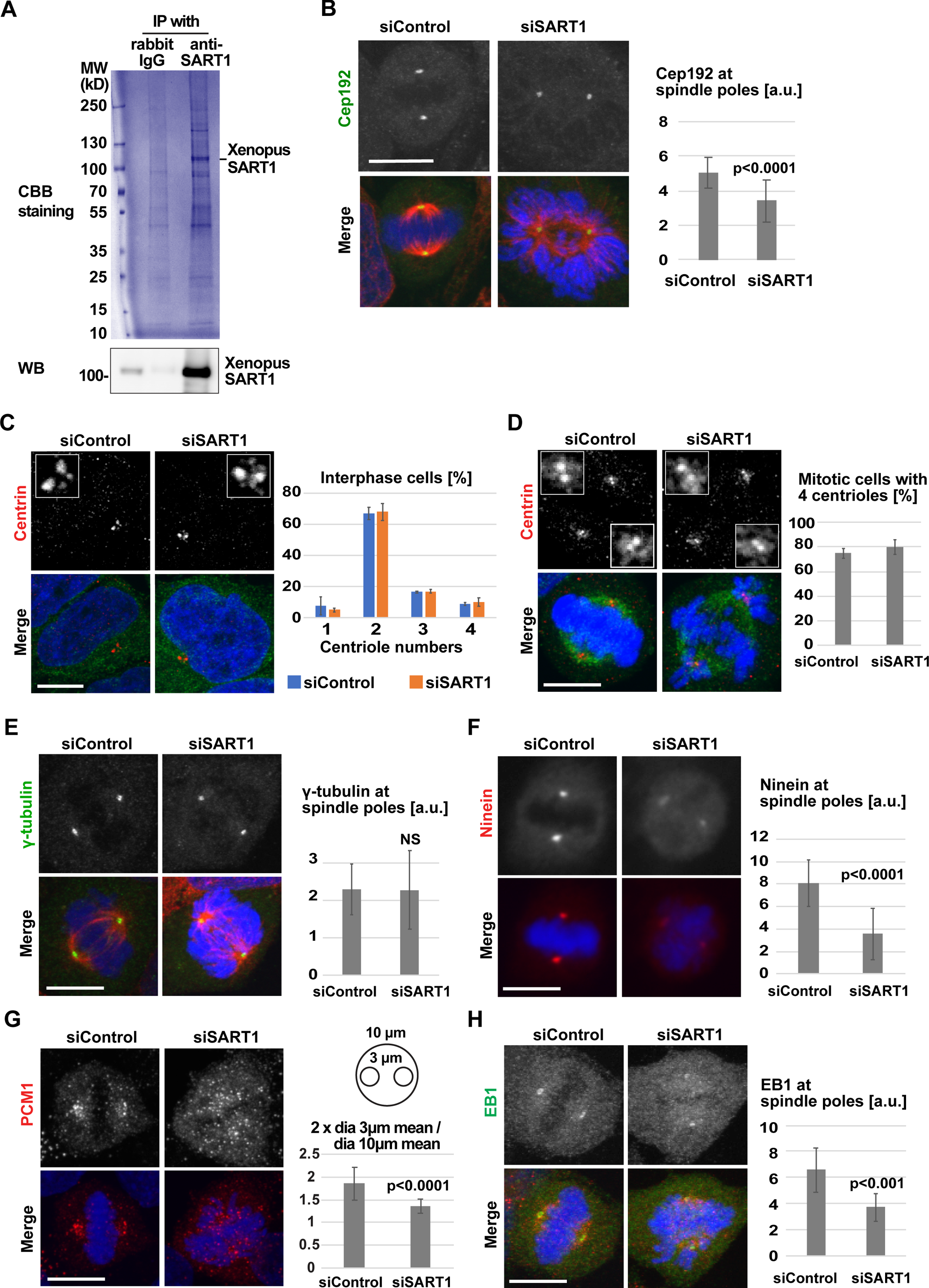
SART1 is required for recruitment of multiple centrosomal proteins. A. Immunoprecipitation (IP) of SART1 from Xenopus egg extract using rabbit polyclonal antibodies raised against Xenopus SART1, analyzed by SDS-PAGE and Coomassie staining. Note the SART1 signal at 110 kDa, confirmed by Western blot. B. Reduction of Cep192 at mitotic centrosomes in SART-downregulated HeLa cells. siRNA-treated cells are stained for Cep192 (green), α-tubulin (red), and DNA (blue). Cep192 intensity at spindle poles were quantified using ImageJ. N = 2 experiments, n= 10 cells per experiment. p value (student’s test, two tailed). C and D. Centrosome duplication is not impeded by SART1 downregulation. siRNA-treated HeLa cells were stained for the centriole marker centrin (red), α-tubulin (green), and DNA (blue). Centriole numbers are counted per cell. N = 2 experiments, n = 50 cells per experiment. C, Interphase. D, Mitosis. E. No change of γ-tubulin at centrosomes. siRNA-treated HeLa cells were stained for γ-tubulin (green), α-tubulin (red), and DNA (blue). γ-tubulin intensity at spindle poles were quantified. N = 2 experiments, n = 10 cells per experiment. F. Reduction of Ninein at centrosomes. siRNA-treated cells were stained for Ninein (red), and DNA (blue). Ninein intensity at spindle poles were quantified. N = 2 experiments, n = 10 cells per experiment. G. Disperse of PCM1 over the mitotic cell upon SART1 downregulation. siRNA-treated cells were stained for PCM1 and DNA. Accumulation of PCM1 at spindle poles was calculated as ratio of PCM1 intensity at the spindle poles (two 3 μm circles) and a larger area around chromatin (10 μm circle). N = 2 experiments, n = 10 cells per experiment. H. EB1 centrosome enrichment is reduced upon SART1 depletion. siRNA-treated cells were stained for EB1 (green), α-tubulin (red), and DNA (blue). EB1 signal at spindle poles was quantified. N = 2 experiments, n = 10 cells per experiment. p values (student’s test, two tailed). NS (not significant). Scale bars, 10 μm.

Following its identification with the highest score among the centrosomal proteins, we examined the localization of Cep192 in SART1-downregulated HeLa cells. Cep192 is an integral component of centrosomes required for both centriole duplication and centrosome maturation (Gomez-Ferreria et al., 2007; Zhu et al., 2008). Immunostaining with a Cep192 antibody showed that Cep192 was partially lost from centrosomes upon SART1 knockdown (Fig. 4B). Because Cep192 is essential for centriole duplication (Zhu et al., 2008), we wondered if SART1 depletion also affects centriole duplication. Immunostaining of the centriole marker centrin showed, however, that the number of centrioles was unaffected by SART1 depletion (Fig. 4C and D). As in the control (Fu et al., 2015), we detected mostly two centriole dots in interphase and four dots (two for each spindle pole) in mitosis, indicating proper centrosome duplication in the absence of SART1 (Fig. 4C and D).

It is known that knockdown of Cep192 prevents centrosome maturation by abolishing the recruitment of PCM proteins containing the MT nucleation factor γ-tubulin (Gomez-Ferreria et al., 2007). SART1-depleted cells did not show reduced amounts of γ-tubulin at centrosomes (Fig. 4E), while the localization of Ninein, PCM1, and EB1 was significantly impaired (Fig. 4F-H). EB1 is a plus-end marker of MTs, but an important centrosome component to anchor MTs (Louie et al., 2004). The reduction of EB1 at centrosomes (Fig. 4H) is consistent with the decreased incorporation of the related protein EB3 in the live cell imaging (Fig. 2A). We asked whether SART1 downregulation changes the amounts of PCM proteins and therefore performed quantitative PCRs and Western blots. The mRNA level of Ninein and PCM1, and the protein level of EB1 were unchanged in the absence of SART1 (Fig. S2A and S5C). One of the quantitative PCR primers was designed over two exons (Table S2), indicating also that mRNA splicing of Ninein and PCM1 was not impaired (Fig. S5C). In agreement with the notion that its protein level was unchanged, PCM1 was dispersed throughout the mitotic cell in the absence of SART1 (Fig. 4G).

Together these data reveal that SART1 depletion does not affect centriole numbers nor γ-tubulin recruitment, but significantly reduces localization of specific PCM components at centrosomes. These results indicate that distinct from Cep192, SART1 is specifically required for recruitment of selective PCM proteins to mitotic centrosomes.

### SART1 promotes spindle bipolarity via its N-terminal microtubule-binding region

To further understand the molecular function of SART1, we used Xenopus egg extracts. In this cell-free system, spindle assembly can be faithfully recapitulated when sperm heads are added to the extracts and progression into interphase and then to mitosis is triggered by cell cycle activation. Our Xenopus SART1 antibody, used for the immunoprecipitation (Fig. 4A), efficiently depleted endogenous SART1 from egg extracts (Fig. 5A). While control-treated (mock) extracts assembled bipolar spindles with chromosomes aligned at the metaphase plate, depletion of SART1 caused defects in spindle assembly (Fig.5B). Although MT polymerization around sperm chromatin was quantitatively similar to the control situation (Fig. 5C), spindle bipolarity was not established, and chromosomes did not align to the spindle center (Fig.5B). These spindle abnormalities were rescued by addition of Xenopus SART1 mRNA to express recombinant SART1 (Fig. 5A, B, and G).

**Figure 5.**
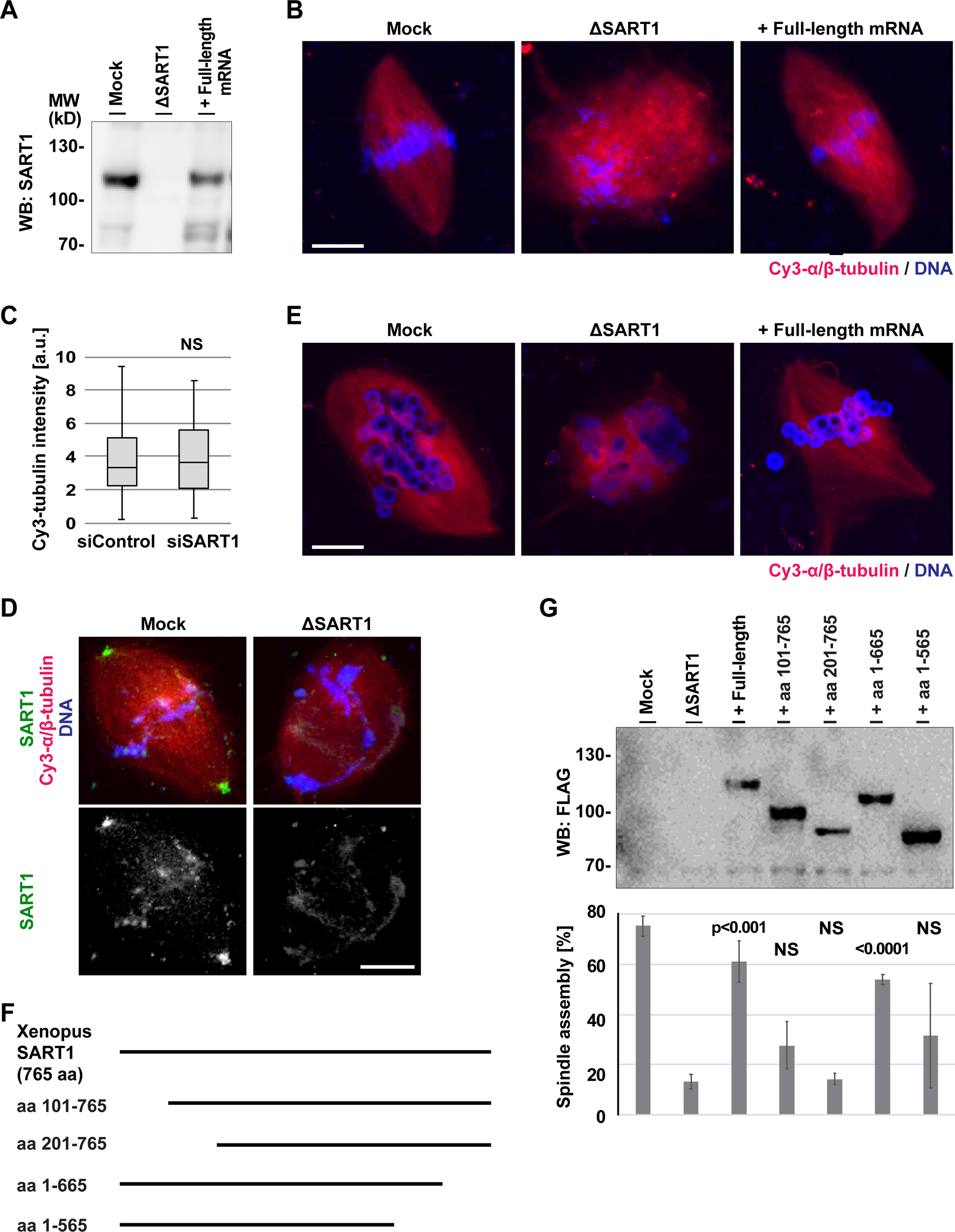
SART1 is required for spindle pole assembly in Xenopus egg extracts and its N-terminus is essential for this function. A. Depletion and addback of SART1, examined by Western blot. Xenopus laevis egg extract was treated with control (mock) or SART1 antibody-coated beads. The resulting extracts were used for the cell cycle reaction in the presence or absence of SART1 mRNA. B. SART1 is required for spindle assembly. Mock- and SART1-depleted extract shown in A were incubated with sperm, Cy3-labeled tubulin (red) in the presence or absence of SART1 full-length mRNA, and cycled to interphase and mitosis. DNA was stained with DAPI. Note that not clear spindle poles were found in depletion and the bipolarity was re-established by expressing recombinant SART. C. SART1 depletion does not affect MT amounts assembled around chromatin. Cy3-tubulin intensity around chromatin was quantified. N = 3 experiments, n = 10 structures per experiment. p values (student’s test, two tailed). NS (not significant). D. SART1 localizes to spindle poles in Xenopus egg extracts. Cycled sperm spindles were assembled as in B, and stained with Xenopus SART1 antibody and DAPI. E. SART1 is essential for spindle assembly around DNA-beads. Mock- and SART1-depleted extract was incubated in DNA-coated beads and Cy3-tubulin in the presence or absence of SART1 mRNA, and cycled for spindle assembly. Note that even the spindles without centrosomes, SART1 is required for spindle pole assembly. F. Xenopus SART1 protein and its deletion mutants constructed. G. The N-terminal, NLS-containing region of SART1 is critical for bipolar spindle assembly. SART1-depleted extract was supplemented with indicated SART1 mRNAs and used for spindle assembly as shown in B. Expression of each exogenous protein was confirmed by Western blot. Frequency of bipolar spindles were counted. N = 3 experiments, n > 50 DNA structures per experiment. p values against ΔSART1 extract (student’s test, two tailed). NS (not significant). Scale bars, 10 μm.

Consistent with our findings in human cells, the Xenopus SART1 antibody stained spindle poles and, importantly, this staining disappeared upon SART1 immunodepletion (Fig. 5D). Using egg extract, spindles were also assembled around DNA-coated beads (Fig. 5E). These artificial spindles do not contain centrioles but still accumulate PCM proteins for spindle pole formation (Heald et al., 1997). Also here, SART1 depletion resulted in spindle bipolarization defects (Fig. 5E) and expression of recombinant SART1 restored bipolarity (Fig. 5E). This demonstrates that SART1 is crucial for spindle bipolarization also in the absence of centrioles.

To understand which parts of SART1 are important for its function in spindle assembly, we constructed N- and C-terminal deletion mutants of the protein (Fig. 5F) and expressed them in SART1-depleted egg extracts (Fig. 5G). Besides full-length SART1, the fragment lacking the last hundred aa at C-terminus (aa1-665) restored bipolar spindle assembly around sperms (Fig. 5G). In contrast, SART1 fragments lacking the N-terminus did not, which suggests that the N-terminal MT-binding domain (Fig. S1D and E) is important for spindle bipolarization. To gain further evidence for this hypothesis, we constructed human SART1 fragments and tested their ability to rescue the spindle abnormality in HeLa cells (Fig. 6A). While full-length SART1 as well as fragments lacking the C-terminus, although less efficiently, restored spindle assembly (Fig. 6B-E), the fragments lacking the N-terminus did not restore the wildtype phenotype. We could not faithfully detect the localization of SART1 fragments in the transfected cells by immunofluorescence, because FLAG antibody nonspecifically stained centrosomes even in non-transfected HeLa cells. Therefore, we prepared lysates from transfected cells and performed a MT sedimentation assays with taxol-stabilized MTs. In agreement with our hypothesis, full-length SART1 and fragments lacking the C-terminus bound to MTs, but fragments lacking the N-terminus did not (Fig. 6C). All these results show that the N-terminus of SART1 is a microtubule-binding domain critical for spindle pole assembly.

**Figure 6.**
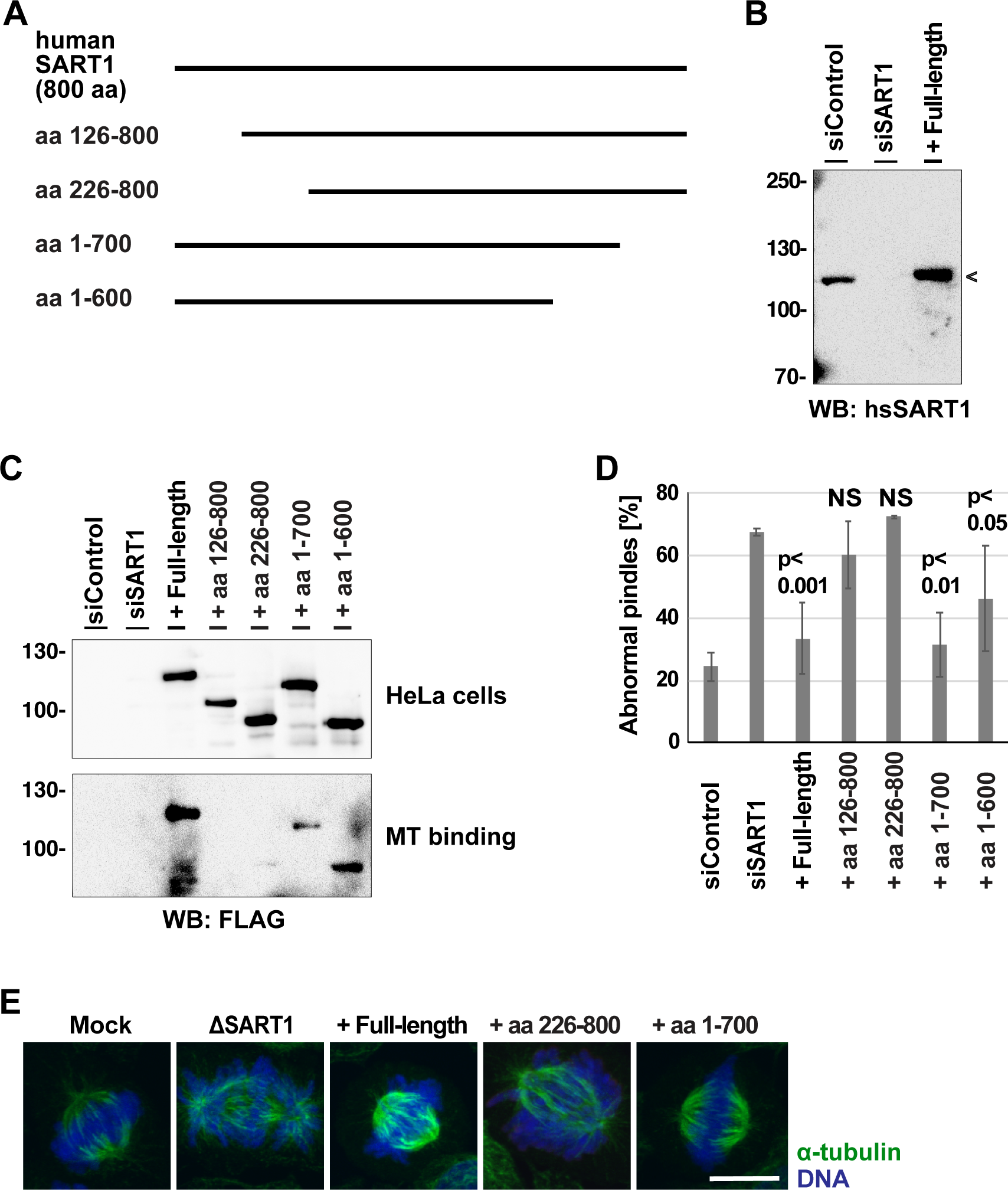
N-terminal, MT-binding region of SART is also important for spindle assembly in human cells. **SART1 depletion causes spindle defects and cell death preferentially in cancer cell lines** A. human SART1 protein and its deletion mutants constructed. B. SART1 mutant lacking N-terminus could not restore spindle assembly defects induced by SART1 silencing. HeLa cells were transfected with full length and deletion mutants of SART1 constructs shown in A. The cells were subsequently treated with control or SART1 siRNAs for 3 days, fixed, stained for α-tubulin and DNA. Protein levels of SART1 were checked by Western blot. Abnormal spindles were counted as in Fig. 1F. Error bars: SD. N = 3 experiments, n > 50 prometaphase and metaphase-like cells per experiment.

## Discussion

SART1 has been previously identified in RNAi screens as a protein required for mitotic progression (Kittler et al., 2004; Neumann et al., 2010) and centrosome biogenesis (Balestra et al., 2013). However, mitotic and centrosomal functions of SART1 have not been characterized in detail and it remained unclear whether SART1 is required for those events directly or indirectly via pre-mRNA splicing of other relevant factors. Here we demonstrate that SART1 acts as a bona fide RanGTP-regulated MAP. SART1 localizes next to centrosomes and is required for centrosome maturation and spindle assembly. Some splicing factors (Cdc5l and the Cdc5l-containing Prp19 complex) have been shown to be similarly important for spindle assembly (Hofmann et al., 2013; Mu et al., 2014). Although these proteins do not localize to specific mitotic structures, their functions on spindle assembly are independent of mRNA splicing (Hofmann et al., 2013). Thus, our results support the idea that multiple splicing factors have moonlighting functions in mitosis (Somma et al., 2020).

Downregulation of SART1 significantly reduces the number of MT plus-ends, MT length and lifetime. Importantly, these defects are specifically detected in mitosis and not in interphase. This can be explained by the fact that SART1 resides inside the nucleus in interphase and only becomes accessible to MTs during mitosis. The mitotic MT defects could be cause of the observed chromosome misalignment, the lack of end-on attachment, reduced inter-kinetochore distance, and checkpoint activation (Foley and Kapoor, 2013). SART1 has been described as one of the pre-mRNA splicing factors required for sister chromatid cohesion (Sundaramoorthy et al., 2014). The reduced inter-kinetochore distance and the results of chromatin spreads show, however, that chromatid cohesion is proper in our SART1-depleted condition.

Upon nuclear envelope breakdown, SART1 localizes around the two centrosomes and further accumulates there in prometaphase. Only when bipolar spindles are established during metaphase, SART1 forms a half circle around centrosomes pointing away from the spindle axis. This unique SART1 localization is dependent on the presence of MTs, suggesting that SART1 is not an intrinsic component of centrosomes like γ-tubulin. We are not aware of another protein with a similar localization pattern as SART1 at the centrosomal periphery. Therefore, we propose to name this novel centrosomal structure as SART1 cap, which is potentially important for centrosomal maturation. How this cap-like structure forms is an interesting question for the future. We envision that pulling forces moves SART1 toward the cell cortex and establishes the structure, similar as outer kinetochores are stretched to opposite directions upon end-on MT attachment (Hara and Fukagawa, 2020; Smith et al., 2016).

We consistently found by immunoprecipitation that SART1 interacts with a number of centrosomal proteins (Cep192, APC, POC1, Cep44, Cep43, Cep152, Cep63, Plk1, SSX2IP, Cep97, Cep85, POC5, CDK11B, Cep70, Centrin 3) in Xenopus egg extracts. We subsequently found that SART1 depletion reduces the accumulation of selective centrosomal proteins at centrosomes, including Cep192, Ninein, PCM1, and EB1, while it did not affect recruitment of the major MT nucleation factor γ-tubulin. Although Cep192 depletion is known to abolish both centriole duplication and centrosome maturation (Gomez-Ferreria et al., 2007; Zhu et al., 2008), SART1 depletion, in which Cep192 was ∼30% reduced, did not affect centriole duplication and the amounts of γ-tubulin at centrosomes. These results suggest that the reduction of other PCM proteins (Ninein, PCM1, and EB1) in SART1-depleted extracts is not caused by the partial reduction of Cep192 but due to a direct, specific effect of the SART1 absence. PCMs proteins recruited by SART1 could promote MT dynamics and, as a result, contribute to chromosome alignment (Foley and Kapoor, 2013; Kumari and Panda, 2018).

While spindles are severely defective in SART1-depleted cells, the amount of spindle MTs seem not to be affected. Indeed, in SART1-depleted egg extract we find similar spindle defects as in human somatic cells with MT amounts not statistically different to control spindles. This phenotype is consistent with the fact that γ-tubulin accumulation at centrosomes is not affected in SART1-depleted human cells. Thus, SART1 is not required for MT nucleation, but essential for spindle bipolarization by recruiting selective PCM proteins. The pole assembly defects are also detected in spindles assembled in vitro around sperm and DNA-coated beads. Importantly, spindles assembled around the DNA-coated beads do not contain centrosomes, but still recruit PCM proteins for spindle assembly (Heald et al., 1997). Taken together, we conclude that SART1 is not required for centrosome de novo formation and MT assembly, but essential for spindle bipolarization as a part of centrosome maturation.

Both in egg extracts and human cells, full-length SART1 rescues spindle assembly defects caused by SART1 depletion, while SART1 fragments lacking the N-terminal domain does not. The SART1 fragments lacking the N-terminal domain does not bind to MTs if transfected into human cells. Consistently, the N-terminal fragment of Xenopus SART1 binds in vitro directly to MTs and bundles pre-existing MTs. Several nuclear proteins bind MTs via their NLS and neighboring residues and regulate MT functions with the other protein domains (Yokoyama, 2016). Indeed, the N-terminus of SART1 contains a predicted NLS (aa 26-46 of Xenopus SART1, aa 31-59 of human SART1 (Kosugi et al., 2009). Together, we conclude that the N-terminus of SART1 contains the MT binding region and is essential for spindle pole formation.

In summary, our results unravel SART1 as a novel RanGTP-regulated MAP that localizes to the outer surface of centrosomes along the spindle axis, which we designate as SART1 cap. SART1 promotes recruitment of specific centrosomal components for spindle pole assembly that are important for MT dynamics and proper chromosome alignment.

## Methods

### Recombinant proteins and antibodies

A cDNA covering the full-length Xenopus laevis SART1 (NM_001086027.1) was in vitro synthesized (GenScript) and subcloned into pET28a (Novagen) with NdeI and XhoI sites. For antibody production, the recombinant Xenopus SART1 protein was expressed in BL21 (DE3) E. coli and solubilized from inclusion bodies with 6 M urea. The protein was purified with Ni-NTA Agarose (Qiagen), dialyzed to PBS containing 6 M urea, and used for immunization in rabbits (Hokkaido System Science). From the antisera, Xenopus SART1 antibody was purified using the antigen column and used for Western blot and immunofluorescence at 1 μg/ml. To increase recombinant Xenopus SART1 solubility, His-Acidic-Target Tag (HisATT), modified from Flag-Acidic-Target Tag (Sangawa et al., 2013), was synthesized (Genscript) and subcloned into the pET28a-xlSART1 plasmid via the NcoI and NdeI sites. The HisATT-SART1 was expressed in BL21 (DE3) and purified with Ni-NTA, and dialyzed to CSF-XB buffer (10 mM K-HEPES, 100 mM KCl, 3 mM MgCl_2_, 0.1 mM CaCl_2_, 50 mM sucrose, and 5 mM EGTA, pH 7.7) containing 10% glycerol and 1 mM DTT, and used for MT sedimentation assay. Xenopus SART1 fragments were amplified by PCR, cloned into pET28a, and prepared as HisATT-SART1. For in vitro translation in egg extracts, Xenopus SART1 and its depletion mutants were amplified by PCR and subcloned into a pCS2+ vector harboring the FLAG tag sequence at the N-terminus. Importin α, importin β, and RanQ69L-GTP were expressed in E. coli and purified with TALON beads (Yokoyama et al., 2014).

The following published and commercial antibodies were used: rabbit (Rb) anti-chTOG for western blot (WB) at 1 μg/ml, Rb anti-α-tubulin (Yokoyama et al., 2014) for immunofluorescence (IF) at 2 μg/ml, anti-γ-tubulin (Barenz et al., 2013) (Rb, IF 1 μg/ml), Ninein serum (Barenz et al., 2013) (Rb, IF 1:500), PCM1 (Barenz et al., 2013) (Rb, IF 1:1000), XCAP-G (Yokoyama et al., 2019) (Rb, WB 1 μg/ml), α-tubulin (mouse (Ms) B-5-1-2; Sigma, IF 1:2000), BubR1 (Ms, BD, IF 1:500), CENP-A (Ms, Enzo, IF 3 μg/ml), Centrin (Ms, Millipore, IF 1:500), Cep192 (Rb, Bethyl, IF 1 μg/ml), CREST (human, Antibody Inc., IF 1:200), EB1 (Ms, BD, IF 1:250), FLAG (Ms M2, Sigma, WB 1:1000), γ-tubulin (Ms GTU-88, Sigma, WB 1:1000, IF 1:500), GAPDH (Ms, Santa Cruz, WB 1:2000), Ndc80 (Ms, GeneTex, IF 1:500), phospho-Histone H3 (Rb, Millipore, IF 1:500), SART1 #1 (Ms, Abcam, IF 3 μg/ml, WB 1 μg/ml), and SART1 #2 (Ms, Santa Cruz, IF 3 μg/ml), and pericentrin (Rb, Abcam, IF 1 μg/ml). Secondary antibodies for immunofluorescence were anti-rabbit IgG conjugated with Alexa Fluor 488 or 568, and anti-mouse IgG conjugated with Alexa Fluor 488 or 568 (above from Life technologies, 1:1000), and anti-human IgG conjugated with CF568 (Biotum, 1:1000). DNA was counterstained with DAPI at 1 μg/ml.

### Sendai viruses (SeVs) and plasmid vectors for rescue experiments in human cells

Human SART1 mutant resistant to siSART1#3 (s228453) was designed by introducing the following silent mutations into the ORF of SART1 cDNA (NM_005146.5): A813G; T819C; T822C; C825T; C828T. The mutant cDNA with N-terminal FLAG tag was synthesized in vitro (Genewiz), and cloned into pSeV/TSΔF (Ban et al., 2011) (Fig. S2E). To reconstruct virus, the plasmid DNA was transfected to LLC-MK2 cells stably expressing F (fusion) protein.

The Human SART1 mutant resistant to siSART1#3 and its deletions mutants (amplified by PCR) were subcloned into pCI-neo Mammalian Expression Vector (Promega).

### MT binding and bundling assays

HeLa nuclear extract (4C Biotech) was diluted to 1 mg/ml with CSF-XB buffer, and centrifuged at 20,000 xg for 10 min at 4°C. The supernatant was incubated with 2 μM pure taxol-stabilized MTs at RT in the presence or absence of recombinant 3 µM importin α/β complex and 5 µM RanQ69L-GTP, a dominant positive mutant of Ran locked in the GTP-bound state, and pelleted at 20,000 g for 10 min 20°C. Microtubule-associated proteins (MAPs) were eluted from the pellet with CSF-XB supplemented with 500 mM NaCl for 5 min. After another centrifugation, the supernatant (eluate) was analyzed by Western blot.

Xenopus CSF egg extracts were diluted 1:3 with CSF-XB buffer. After centrifugation at 20,000 xg for 10 min at 4°C, the supernatant was incubated at RT in the presence or absence of 4 µM taxol-stabilized microtubules for 15 min. The samples were centrifuged at 20,000 xg for 10 min at 20°C, and pellets were incubated with CSF-XB supplemented with 500 mM NaCl for 5 min, and centrifuged again. The resulting supernatant (eluate) and pellet were analyzed by Western blot.

0.1 µM recombinant SART1 was incubated with 2 µM taxol-stabilized MTs for 15 min at RT, and centrifuged at 20,000 xg for 10 min at 20°C. The Supernatant and pellet were analyzed by Coomassie staining or Western blot. The assay was also done in the presence or absence of recombinant 3 μM importin α, 3 μM importin β, and 5 μM RanQ69L-GTP.

To examine MT bundling activity, 0.3 μM Cy3-labeled MTs was incubated with ∼1 µM recombinant Xenopus SART1 aa1-400 in 10 µl BRB80 (80 mM PIPES, 1 mM MgCl_2_, 1 mM EGTA, pH 6.8) buffer for 10 min at RT. The samples were squashed between slides and coverslips with fixative. Images were acquired by using an Olympus Fluoview FV1000 confocal microscope

To examine the MT binding ability of SART1 fragments expressed in HeLa cells, cells were lysed in RIPA buffer (50 mM Tris pH 8.0, 150 mM NaCl, 50 mM sodium orthovanadate, 1% v/v NP40, 0.1 mM PMSF) supplemented with complete protease inhibitor cocktail (Roche). The cell extract (200 μg) was incubated with 2 µM taxol-stabilized MTs for 15 min at RT, and centrifuged at 20,000 xg for 10 min at 20°C. The pellet containing MTs and MAPs was analyzed by Western blot.

### Cell Culture, transfection, immunofluorescence, and microscope

HeLa and U2OS cell lines were cultured in Dulbecco’s modified Eagle’s medium (DMEM) supplemented with 2 mM GlutaMAX, 10% FBS and 500 units/ml penicillin-streptomycin (all from Gibco). The knockdown experiments were performed with the following siRNA oligonucleotides: siSART1#1 (s17343), 5′- GGCUCAACAUGAAGCAGAAtt-3′, siSART1#2 (s17345), 5′- CCCAAUACAGCUUACCGUAtt-3′, siSART1#3 (s228453). AllStars siRNA (from Qiagen) was used as negative control. HeLa and U2OS cells were grown on 12 mm round coverslips (Marienfeld) and transfected with 10 nM of each siRNA using Lipofectamine RNAiMAX (Invitrogen) according to the manufacturer’s instructions. After 72 h, cells were fixed with Mildform 10N (Wako) at RT for 15 min and stored at 4°C. For immunostaining with BubRI and centrin antibodies, cells were fixed with cold methanol, instead of Mildform, at −20°C for 10 min. Unless otherwise stated, siSART1#3 was used for the functional analyses of SART1, including the rescue experiments.

For rescue experiments, HeLa cells were infected with SeVs at MOI3. After 8 h incubation, medium was replaced to fresh one containing 20 nM siRNAs, and cells were cultured for additional 72 h. Alternatively, HeLa cells were transfected with pCI-neo-hsSART1 full-length and ideletion mutants with 1 ng/μl using ViaFect Transfection Reagent (Promega). After 4 h incubation, the siRNA and Lipofectamine mixture was added to the cells and incubated for additional 72 h.

When indicated, HeLa cells were treated with 20 μM nocodazole for 10 min, and fixed with Mildform.

For immunofluorescence staining, fixed cells were incubated with blocking buffer (PBS + 2% BSA + 0.1% Triton-X100) at RT for 30 min, and then incubated with the primary antibodies in the blocking buffer at 4°C overnight. Cells were washed with PBS, incubated with the secondary antibodies and DAPI at RT for 30 min, washed again with PBS, and mounted with Mowiol 4-88 (Calbiochem). To quantify abnormal spindles objectively, we considered prometaphase cells, which may be going to progress to metaphase, as abnormal cells. Thus, the abnormal spindles contain normal prometaphase cells.

Fluorescence images were acquired by using an Olympus Fluoview FV1000 confocal microscope equipped with a UPlanSApo 60x/1.35 Oil objective at 0.5 μm Z steps. Maximum intensity projections were obtained using ImageJ (NIH). Confocal slices are used to detect kinetochore-MT attachment and sister kinetochore pairs. Interkinetochore distance was measured using ImageJ. Maximum projected images are used to quantify fluorescent intensity using ImageJ. Accumulation of PCM1 at spindles poles was calculated as ratio of PCM1 signal intensity at poles (two 3 μm circles) and larger area around chromatin (10 μm circle) (Fig. 4).

### Live cell imaging experiments

HeLa cells expressing H2B–mCherry were transfected with 20 μM siRNA oligonucleotides in 8-well µ-slide chambers (Ibidi) and, after 30 h, were imaged for 48 h in a Axioobserver Z1 (Zeiss) equipped with a heating and CO_2_ incubation system (Ibidi), Colobri LED illumination (Zeiss), a CCD camera (AxioCamMR3; Zeiss), and a Plan-Apochromat 10× NA 0.45 M27 objective. Single position files were converted into image sequences with the AxioVision software (LE64; V4.9.1.0; Zeiss). Afterwards, segmentation, annotation, classification of cells was performed using the CellCognition software (http://www.cellcognition.org/software/cecoganalyzer) (Held et al., 2010; Yokoyama et al., 2019). More than 100 cell mitotic trajectories per condition were used for quantification and analysis with Microsoft excel and GraphPad Prism.

HeLa cells expressing GFP-tubulin and H2B–mCherry (Held *et al*., 2010), a kind gift from Daniel Gerlich (IMBA, Vienna), were transfected with 20 μM siRNA oligonucleotides in 8-well µ-slide chambers (Ibidi) and, after 30h, were imaged with a Ti2 Eclipse microscope (Nikon, Melville, NY, U.S.A.) equipped with an X-light spinning disk, a LED light engine SpectraX (Lumecor, Beaverton, OR, USA), GFP/mCherry filter sets and a Plan-Apochromat 40× NA 0.95 objective. A software-based autofocus module from the AR-Elements software (Nikon) was used to perform confocal fluorescence imaging of the single best-in-focus optical section of cells undergoing mitosis every 3 min. Image galleries from mitotic cell trajectories were assembled using FiJi and mounted for figures using Inkscape (Free Software Foundation, Inc. Boston, USA).

To image MT plus ends, HeLa EGFP–EB3-expressing cells (Sironi et al., 2011) were seeded into ibidi 8-well chambers (ibidi GmbH, Germany). Imaging was performed on a Zeiss LSM 880 confocal microscope with a 63/1.4 NA oil-immersion objective in home-made 37°C microscope incubator using medium without Phenol Red (Thermo). Time-lapse images were acquired with a 600-ms exposure at a temporal resolution of 1000 ms for 60 s. Cropping of single frames was performed with in ImageJ using a Gaussian blur filter of radius 2 and analyzed with a multiple particle tracking software (Sironi et al., 2011; Tegha-Dunghu et al., 2014). After processing raw data with ImageJ, we used a MATLAB-based program to detect and track the tips of polymerizing MTs. Tracking algorithms are as described previously (Sironi et al., 2011; Tegha-Dunghu et al., 2014).

### Flow cytometry

3 day after siRNA transfection, cells were stripped with trypsin and stained with a FITC Annexin V Apoptosis Detection kit (BD) following manufacture’s instruction, and analyzed using a CytoFLEX flow cytometer (Beckman Coulter) and FlowJo (FlowJo, LLC).

### Chromosome spreads

HeLa cells were transfected with the indicated siRNAs and grown for 72 h. To enrich mitotic cells, 330 nM nocodazole was added and incubated for 4 h. Cells were stripped by trypsin, washed with PBS, incubated with 75 mM KCl at RT for 8 min. An equal volume of Carnoy’s fixative (methanol: acetic acid = 3:1) was added and centrifuged to remove the supernatant. The Carnoy wash was repeated 2 more times. Cells were resuspended in Caynol, spotted on heated slide glass drop by drop, and stained with DAPI for microscopy. Images were taken using an Olympus Fluoview FV1000 confocal microscope

### Quantitative real-time PCR

3 day after siRNA transfection, total RNAs were isolated using PureLink RNA Mini Kit (Ambion) and were subjected to PureLink DNase (Invitrogen) treatment. Reverse transcription and quantitative real-time PCR were conducted using TaqMan RNA-to-Ct 1-step Kit and 7500 Fast Real-Time PCR System (both from Applied Biosystems) with the indicated primer and probe sets (Table S2). To amplify only spliced mRNAs, one of the primers or probes was designed on exon-intron junction. Values were normalized to those of GAPDH.

### Xenopus egg extracts and cell-free assay

Cytostatic factor-arrested M-phase Xenopus laevis egg extracts (CSF extracts) were prepared as described (Hannak and Heald, 2006). In short, Xenopus eggs were dejellinated by cysteine treatment, washed with CSF-XB buffer (10 mM K-HEPES, 100 mM KCl, 3 mM MgCl_2_, 0.1 mM CaCl_2_, 50 mM sucrose, and 5 mM EGTA, pH 7.7), and crushed by centrifugation at 20,000 g for 20 min in a SW55 Ti rotor (Beckman) at 16°C. The straw-colored middle layer was recovered as a CSF extract. Endogenous SART1 was depleted from CSF extracts by 2 rounds of incubation with 60% (vol/vol) Dynabeads Protein A (Invitrogen) coupled with Xenopus SART1 antibodies.

For spindle assembly in cycled extract (Hannak and Heald, 2006), CSF extract was supplemented with demembraned sperm or DNA-coated beads, and incubated with Cy3-labeled tubulin, and 0.4 mM CaCl_2_ at 20°C for 90 min to allow cell cycle progression into interphase. Samples were cycled to mitosis by addition a fresh CSF extract and incubation at 20°C for 80 min. The spindle structures were spun down to coverslips, fixed with cold methanol, stained with DAPI in PBS, and mounted as described (Hannak and Heald, 2006). For rescue experiments, mRNAs encoding Xenopus SART1 full-length and deletion mutants were prepared from the pCS2+ vectors using mMESSAGE mMachine SP6 kit (Invitrogen). The mRNAs were added to the depleted extract at the beginning of the cell-cycle reactions.

MT density around sperm was quantified using ImageJ. To detect the localization of SART1, the fixed spindles on coverslips were stained with the Xenopus SART1 antibody and subsequently with anti-rabbit IgG conjugated with Alexa Fluor 488 and DAPI.

### Immunoprecipitation of SART1 and mass spectrometry

Dynabeads Protein A were coupled with rabbit IgG or the Xenopus SART1 antibody following the manufacturer’s instruction. The antibody beads were cross-linked with dimethyl pimelimidate. Each bead sample (600 μl slurry) was incubated with Xenopus CSF egg extracts (1000 μl) at 4°C for 60 min, washed twice with CSF-XB and twice with CSF-XB containing 0.5 M KCl and 0.1% Triton X-100. The immunoprecipitates were resuspended in SDS sample buffer and resolved by SDS-PAGE for Coomassie stain or immunoblot.

For shotgun mass spectrometry, the immunoprecipitates (600 μl bead slurry) were eluted from the beads with 0.1 M triethylamine (pH 11.5, 60 μl), and neutralized by addition of final 0.1 M Tris pH 6.8. The proteins were trypsin digested and analyzed with a Q-Exactive mass spectrometer (Thermo Fisher Scientific) at the CoMIT in Osaka University.

Tandem mass spectra were extracted for database searching. Charge state deconvolution and deisotoping were not performed. All MS/MS samples were analyzed using Mascot (Matrix Science, London, UK; version 2.5.1). Mascot was set up to search the Ani_Uni_Xenopus_200130 database (unknown version, 57926 entries) assuming the digestion enzyme trypsin. Mascot was searched with a fragment ion mass tolerance of 0.020 Da and a parent ion tolerance of 10.0 PPM. O+18 of pyrrolysine and carbamidomethyl of cysteine were specified in Mascot as fixed modifications. Deamidated of asparagine and glutamine and oxidation of methionine were specified in Mascot as variable modifications.

### Statistical analyses

For live cell imaging, data were tested for normality by D’Agostino and Pearson omnibus test. For normal distributions, ANOVA test and subsequent Dunnett multiple comparisons test were applied. When normal distributions could not be assumed, statistical significance at alpha = 0.001 was determined using a Kruskal–Wallis test followed by Dunn s multiple comparisons test.

For other fixed cell and egg extract experiments, student’s t-test was performed using Microsoft Excel with two-tail distribution and two-sample equal variance.

## Acknowledgements

We thank Takahiro Maeda, Takayuki Kitogo, Yohei Morita, Yasuji Ueda, Tsuyoshi Tokusumi and Kumiko Saeki for technical advices. We also thank Umihito Nakagawa for conducting mass spectrometry, Daniel Gerlich for providing human cell lines, and Junichi Takagi for sequence information of the Flag-Acidic-Target Tag. This study was supported by JSPS KAKENHI Grant Number JP16K21749 to H.Y.

## Author contributions

H.Y. conceptualized and supervised the project, and performed in vitro, human cell and frog egg extract experiments. H.Y. also analyzed mass spectrometry data. D.M.A. set up, performed and analysed the live cell imaging screening. K.T. prepared DNA constructs, reconstituted SeVs. D.M.A., Z.C., A.S., W.A. and O.J.G. conducted additional live cell imaging and analysis. T.F. performed RNAi/rescue experiments. J.M. prepared Xenopus SART1 antibody. Y.H. performed flowcytometry and qPCR, and maintained a frog facility. H.Y. wrote the paper and H.Y., D.M.A., W.A., and O.J.G. revised it.

