## Supplementary figures and images for "SART1 localizes to spindle poles forming a SART1 cap and promotes spindle pole assembly"

### Figures S1-5

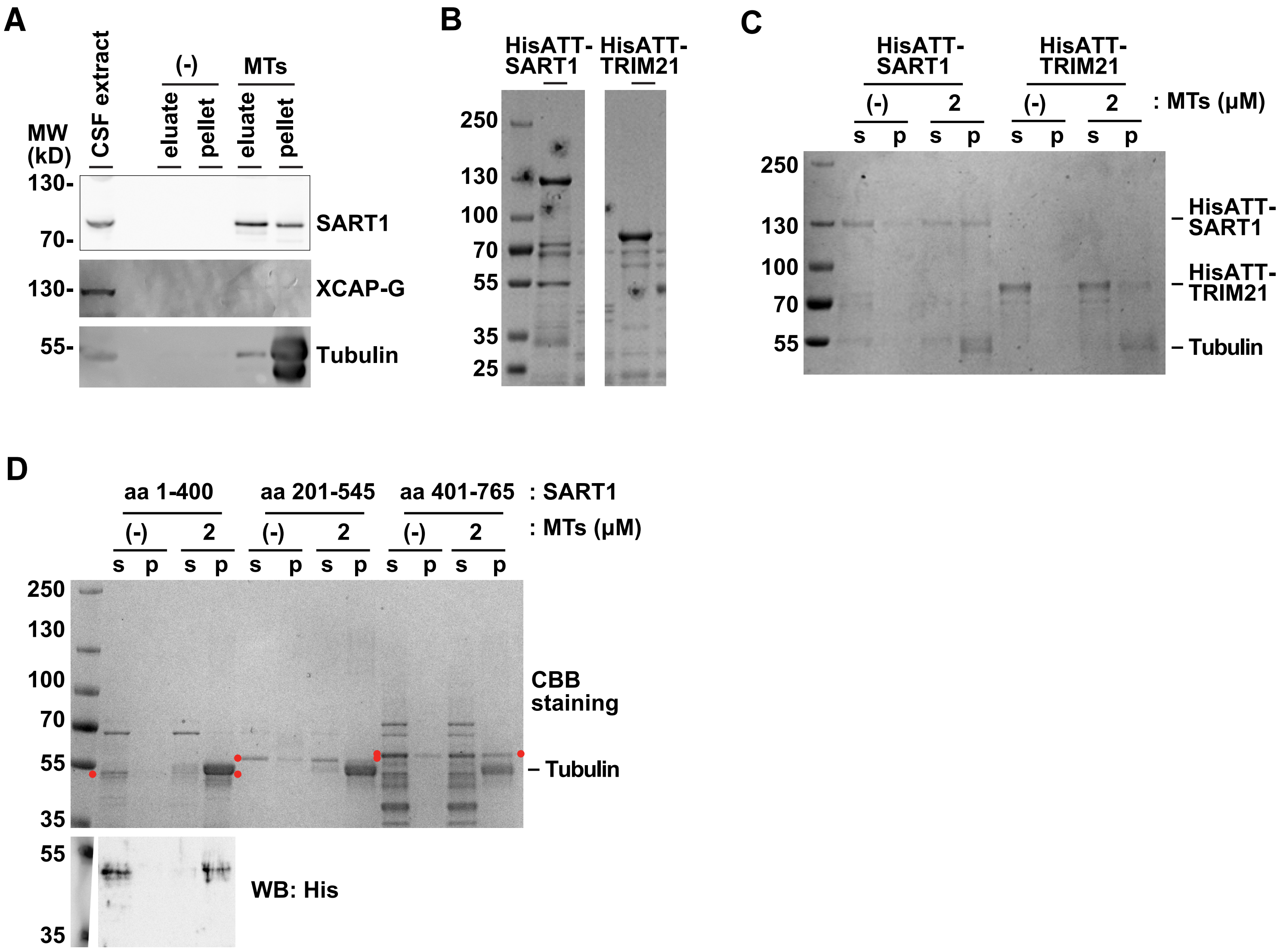

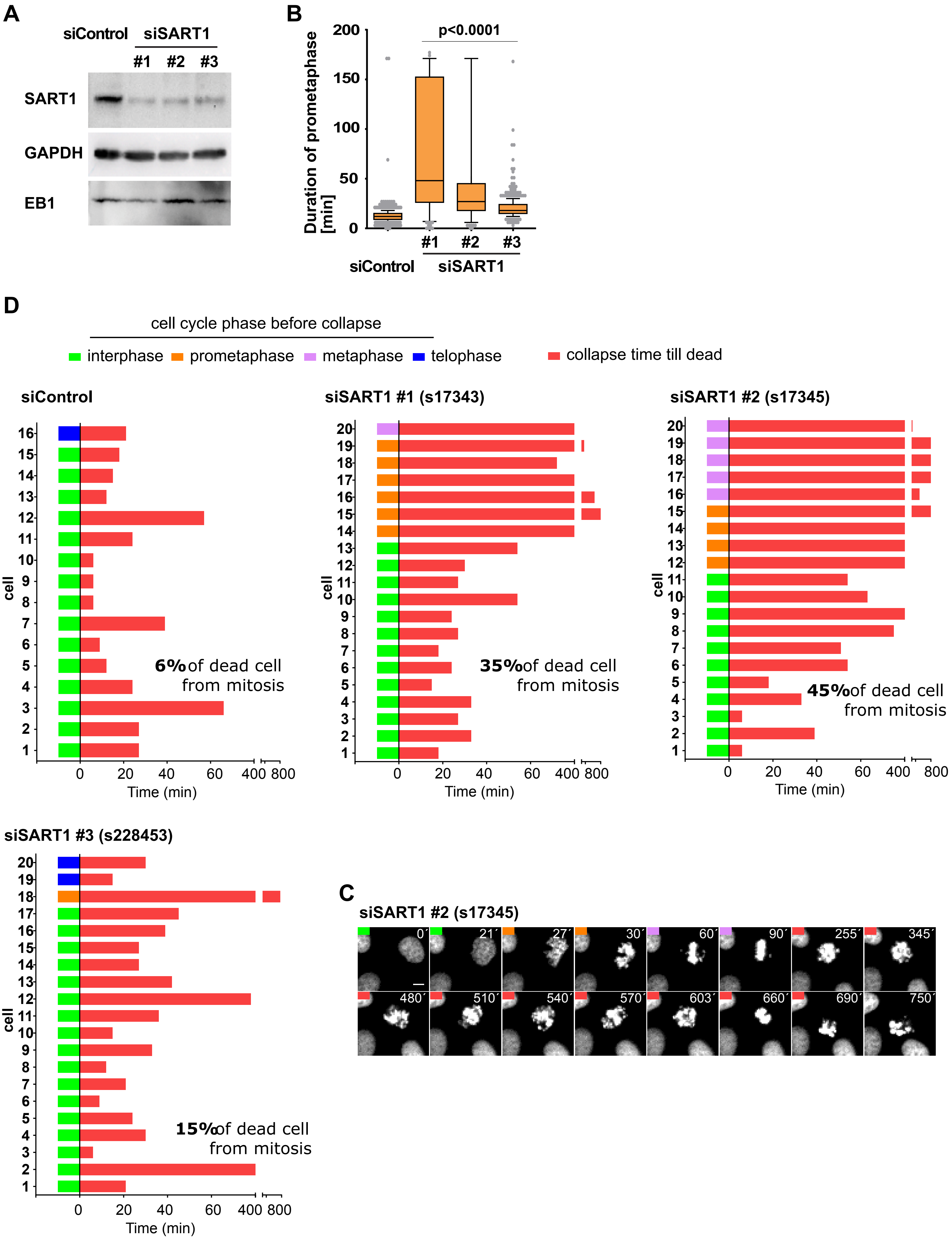

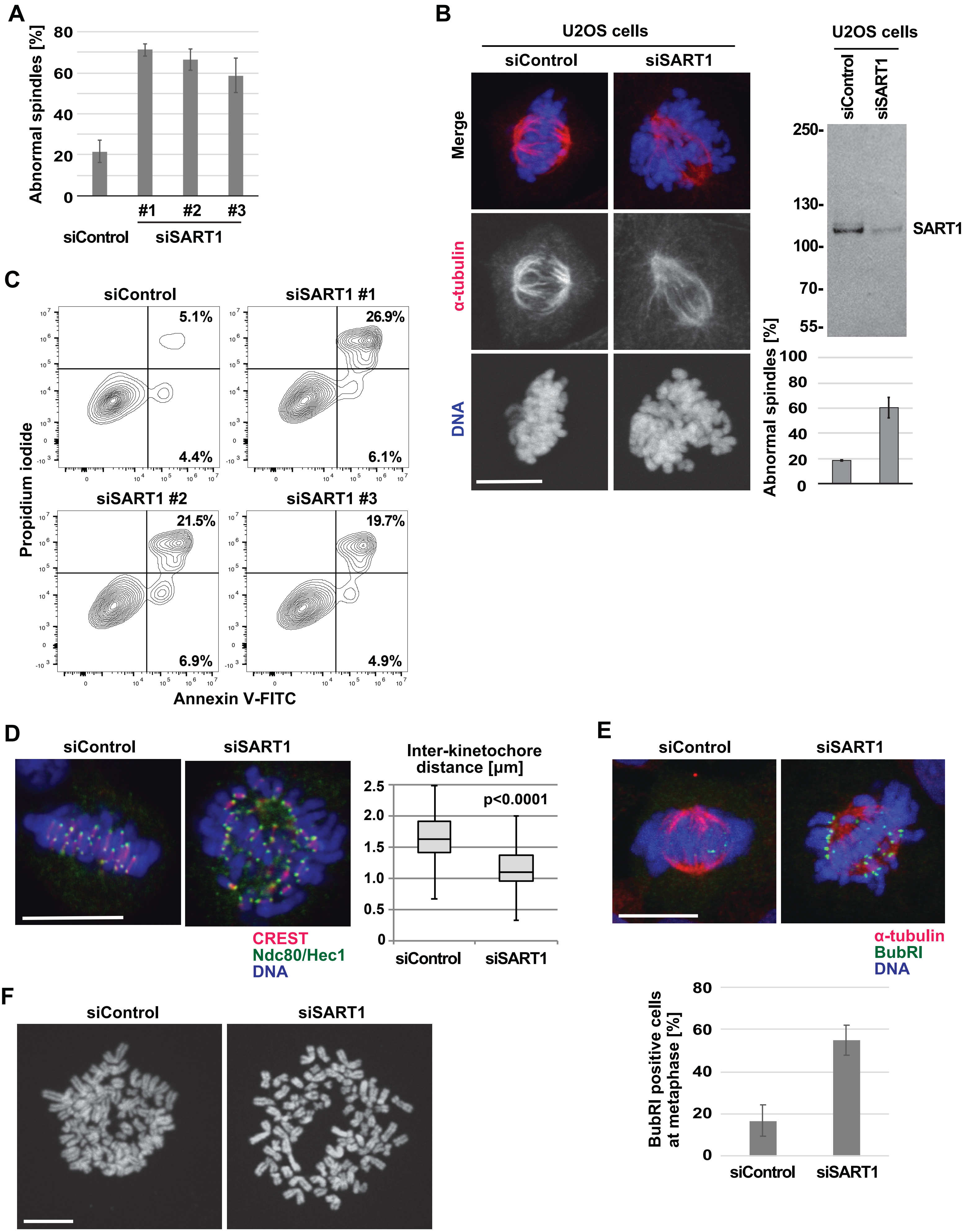

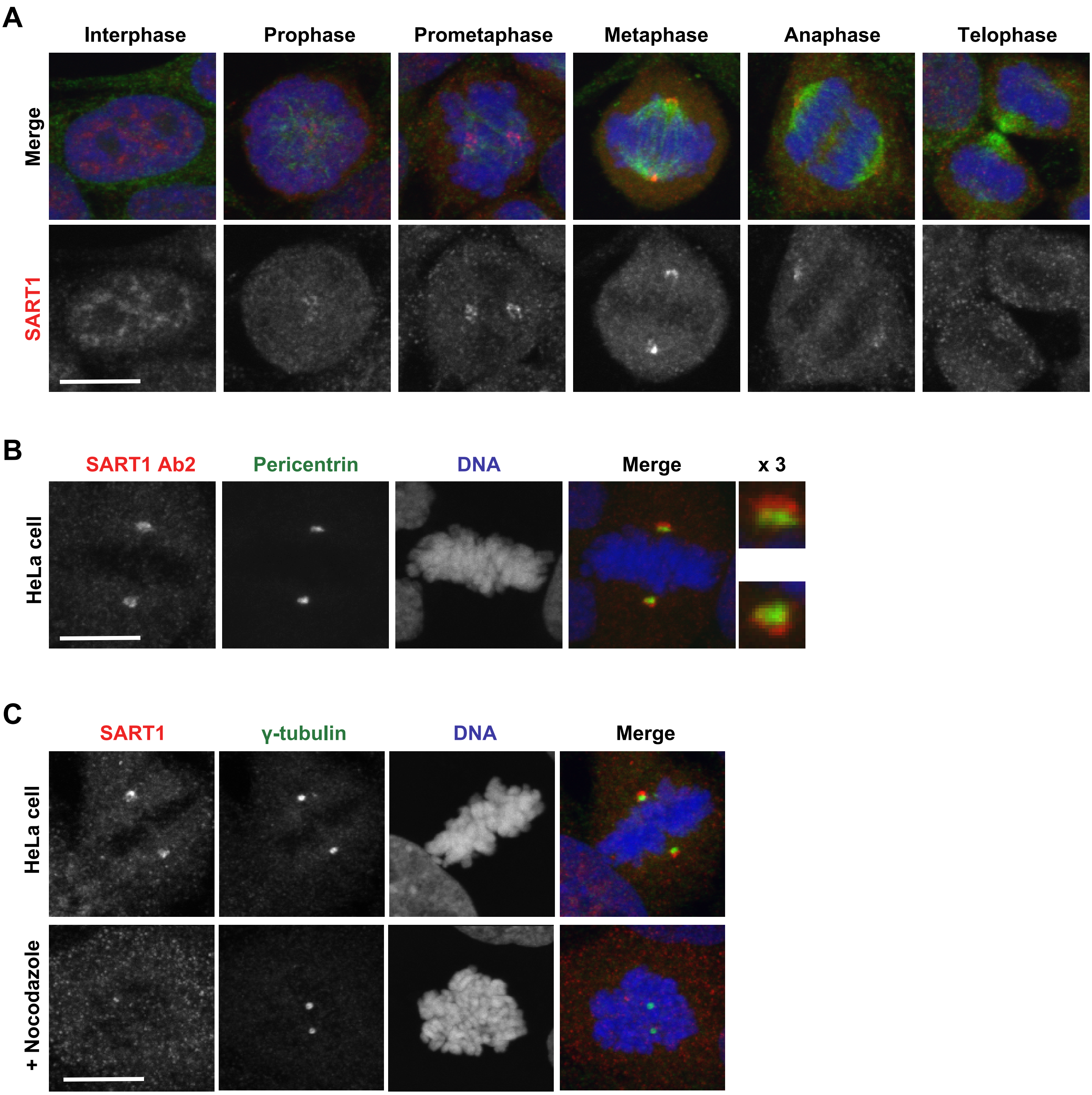

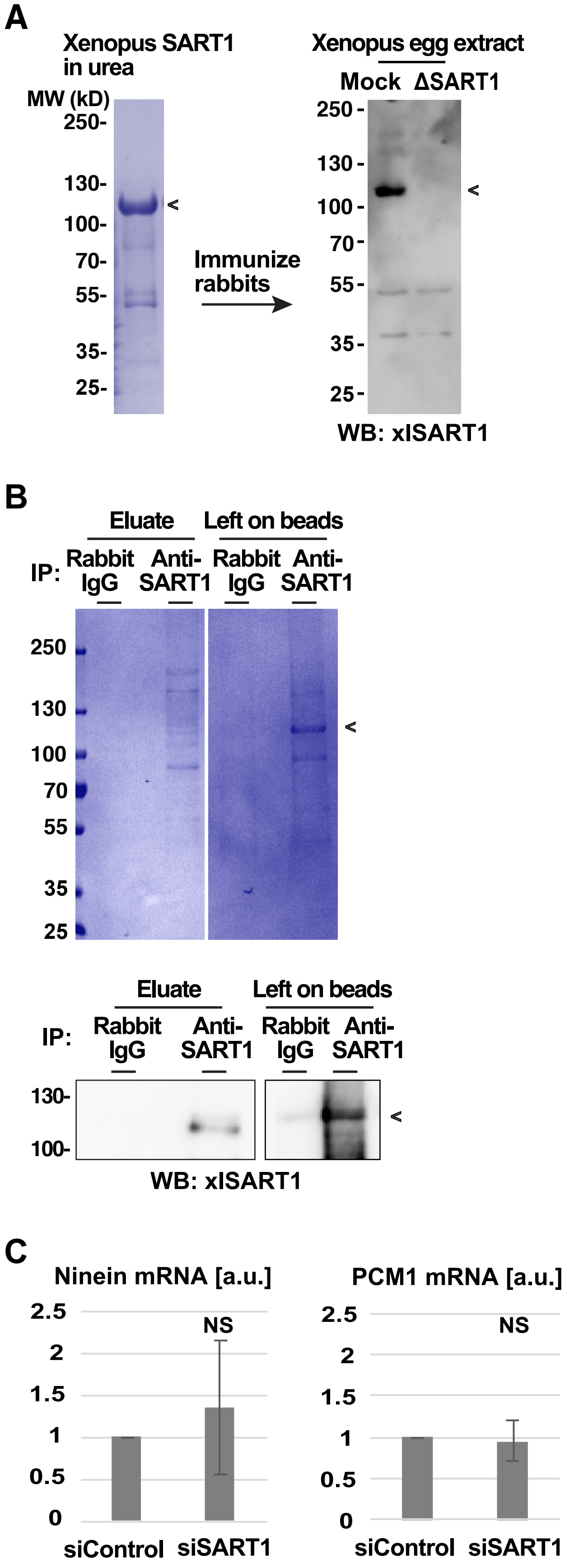
