## Supplementary material for "SART1 localizes to spindle poles forming a SART1 cap and promotes spindle pole assembly": Figures S1-5 legends

**Supplemental Figure legends**

**Figure S1. SART1 is a microtubule-associated protein that binds via its N-terminus.**

A. Endogenous Xenopus SART1 binds to MTs. Xenopus CSF egg extract was incubated with taxol-stabilized MTs. MTs and MAPs were spun down. MAPs are eluted from MTs with high salt and separated by centrifugation. The eluate and MT pellet after elution were immunoblotted for SART1 and XCAP-G as a negative control.

B. His Acidic-Target Tag (HisATT)-fused SART1 expressed in bacteria and subsequently purified with Ni-beads. Note that without the HisATT fusion, SART1 could not be solubilized. As a negative control HisATT-TRIM21 was prepared following the same procedure as for SART1.

C. SART1 interact with MTs in vitro. HisATT- SART1 and HisATT-TRIM21 were incubated with the taxol-stabilized MTs. After centrifugation, supernatant (s) and pellet (p) were analyzed by SDS-PAGE and Coomassie staining. Note that half of HisATT-SART1 was co-sedimented in p, while HisATT-TRIM21 was not.

D. The N-terminal fragment of SART1 binds to MTs directly. Xenopus SART1 fragments were incubated with taxol-stabilized MTs. After centrifugation, supernatant (s) and pellet (p) were analyzed.

**Figure S2. SART1 downregulation in HeLa cells causes prometaphase delay and subsequent cell death.**

A. Western blot of HeLa cells treated for 3 d with 10 μM of three different siRNAs targeting SART1.

B. SART1 downregulation extends prometaphase. Duration of prometaphase was quantified for more than 100 mitotic HeLa cells expressing H2B-mCherry per condition. Statistical significance at alpha = 0.0001 was determined using a Kruskal–Wallis test followed by Dunn´s multiple comparisons test.

C. Example of a time lapse sequence from A showing the collapse and death of a mitotic HeLa cells expressing H2B-mCherry transfected with the indicated siRNA. Scale bar, 10 μm.

D. HeLa cells expressing H2B-mCherry were transfected with 20 μM of the indicated siRNA oligos and after 30h imaged for 70h. Up to 20 randomly chosen dead cells, determined by chromatin fragmentation, per condition were manually tracked back in order to determine the last healthy cell cycle phase (see color legend). Red bars indicate the period since chromatin collapse starts until the first clear chromatin fragmentation appears.

**Figure S3. Similar spindle defects observed with three independent SART1 siRNAs and in also U2OS cells.**

A. Frequency of abnormal spindles in HeLa cells treated as in Fig. 1D. Spindles were stained for α-tubulin and DNA. Error bars: SD. N = 2 experiments, n > 50 prometaphase and metaphase-like cells per experiment.

B. U2OS cells show spindle defects upon SART1 downregulation. BJ, RPE1, and U2OS cells treated with siRNAs as in A were stained for α-tubulin and DNA. Abnormal spindle structures were counted as in Fig. 1E. N = 3 experiments, n > 50 prometaphase and metaphase-like cells.

C. siRNA-transfected HeLa cells as in A were stained with Annexin V and Propidium iodide (PI) and analyzed by flow cytometry. Annexin V positive cells, shown with %, indicate dead cells. Cells in early apoptosis are Annexin V positive and PI negative, and cells in late apoptosis or already dead are both Annexin V and PI positive.

D. siRNA-treated HeLa cells were stained as indicated. Inter-kinetochore distance was measured based on Ndc80 dots along the CREST rods. n > 50 kinetochore pairs. p values (student’s test, two tailed).

E. The spindle checkpoint is active/remains unsatisfied in the absence of SART1. siRNA-transfected HeLa cells were stained for a spindle checkpoint protein BubR1, α-tubulin, and DNA.

F. Chromosome spreads prepared from siRNA-transfected HeLa cells (as in A).

Scale bars, 10 μm.

**Figure S4. Upon mitotic onset, SART1 localizes first around centrosomes and later on the distal side of centrosomes along the spindle axis.**

A. HeLa cells were fixed, and stained for SART1 (red), MTs (green), and DNA (blue). Mitotic stages were classified based on MTs and DNA morphology.

B. HeLa cells were fixed, and stained with an SART1 mouse monoclonal antibody (Ab2, Santa Cruz), pericentrin rabbit polyclonal antibody, and DAPI.

C. In contract to SART1, γ-tubulin centrosomal staining is unaffected by nocodazole treatment. HeLa cells were incubated in nocodazole for 10 min, fixed, and stained for SART1, γ-tubulin, and DNA.

Scale bars, 10 μm

**Figure S5. Immunoprecipitation of SART1 from Xenopus egg extract.**

A. Production of Xenopus SART1 antibody. Full length Xenopus laevis SART1 was expressed in bacteria, purified with Ni-NTA under denaturing condition, and used for antibody production in rabbits. The purified antibody recognizes a SART1 in egg extracts by Western blot. The specificity was confirmed by disappearance of the respective signal after SART1 depletion form egg extracts using SART1 antibody-immobilized beads.

B. Immunoprecipitation (IP) of SART1 was conducted from Xenopus egg extract using antibody beads. Interacting proteins were eluted with triethylamine at pH11.5. The eluate and proteins left on the beads were analyzed by SDS-PAGE and Coomassie staining as well as Western blotting.

C. Ninein and PCM1 mRNA levels were unchanged upon SART1 downregulation. Quantitative real-time PCR was performed and gene expression was normalized by GAPDH levels. Error bars: SD. N = 4 experiments. p value (student’s test, two tailed). NS (not significant).
