## Supplementary material for "SART1 localizes to spindle poles forming a SART1 cap and promotes spindle pole assembly": Table S2

**Table S2. Target sequences of primers and probes used for quantitative real-time PCR**

| Gene name | Primer name | Target sequences (5'-3') |
| --- | --- | --- |
| Ninein | Probe | TCATACTCCTCACTGCGTTGCGTCTTCCA |
|  | Forward | GACGGTGATTGAGCCACTGG |
|  | Reverse^a^ | GTTCCAAAACCTTAACTGGCCTTC |
| PCM1 | Probe^a^ | TGTGATACTGACGCCAGATAAGCTACCTGC |
|  | Forward | TGCAGTGATGGATGATTCTGTTG |
|  | Reverse | CGCTGAATTAAGTCATTCAATTCTTC |

^a^ The primer designed over two exons
